# Loss of PKCθ-GADD45a axis drives triple-negative breast cancer cells into p53-independent senescence

**DOI:** 10.1101/2024.10.10.617550

**Authors:** Amandine Nicolle, Ye Zhang, Valérie Choesmel-Cadamuro, Xiaobo Wang, Karine Belguise

## Abstract

**Background:** PKCθ is a serine/threonine kinase that is well known for its role in the immune system. However, increasing evidence implicates PKCθ in the pathology of breast cancer. PKCθ is highly expressed in triple negative breast cancer (TNBC) cells in which it controls cell migration and invasion, while its implication in cell proliferation remains poorly understood.

**Methods:** To determine the function of PKCθ in cell proliferation, siRNAs were used to modulate the expression of PKCθ in TNBC cells (MDA-MB-231, MDA-MB-436, HCC1937) and cell growth was examined by clonogenic and EdU assays. β-galactosidase assay, RT-qPCR and western blot were used to characterize the senescence features. PCR microarrays and rescue experiments were conducted to investigate the underlying mechanism.

**Results:** We show that PKCθ inhibition leads to a growth arrest in TNBC cells harboring a p53 loss-of-function mutation. This p53-independent growth arrest is accompanied by an increased activity of senescence-associated β-galactosidase, the presence of a senescence-associated secretory phenotype, and the striking expression change of various genes implicated in cell proliferation and senescence. Thus, our data show that PKCθ silencing drives TNBC cells into a senescence-like phenotype. Mechanistically, we demonstrate that p27 is the main CDK inhibitor controlling the PKCθ loss-induced senescence. The accumulation of p27 is due to a surprising strong reduction in GADD45a expression. Indeed, similar to PKCθ silencing phenotype, GADD45a knockdown drives TNBC cells into a senescence-like phenotype.

**Conclusions:** Altogether, our study highlights that the loss of PKCθ-GADD45a axis triggers a p27-dependent senescence response in TNBC cells and further supports strategies targeting PKCθ as treatment for this type of aggressive breast cancer.

## Background

Triple-negative breast cancer (TNBC), which lacks estrogen receptor (ER) and progesterone receptor (PR) expression as well as human epidermal growth factor receptor 2 (HER2) amplification, constitutes 10%–20% of all breast cancers. TNBC exhibits aggressive characteristics such as poor differentiation, a higher rate of proliferation, and increased metastatic capability. Currently, no clinically relevant biomarkers can be used to guide the use of targeted therapies for TNBC and the only treatment available is the toxic, non-specific chemotherapy drugs^1, 2^. Consequently, there is an urgent need to identify more effective, less toxic therapeutic strategies, and thus further investigations of the biology of TNBC are required to identify the signaling pathways that drive the strong aggressiveness of TNBC.

Senescence is a major anti-cancer defense, and it is characterized by a proliferative arrest that prevents the proliferation of damaged cells. This proliferative arrest is induced by various stresses such as DNA damage, oxidative stress, oncogene activation, mitochondrial dysfunction, or chemotherapy. The two signaling pathways that predominantly govern the senescence program are the p53/p21(Cip1) and p16(INK4a)/Rb signaling axes^3–5^. Lately, p27(Kip1) has begun to emerge as another key effector of cellular senescence^6, 7^. There are several features for senescent cells. Firstly, senescent cells display striking changes in gene expression such as changes in cell cycle regulators. Secondly, the cell arrest is accompanied by increased senescence-associated β-galactosidase (SA-β-gal) activity and by the appearance of senescence-associated heterochromatin foci (SAHF). Finally, senescent cells secrete a plethora of molecules such as inflammatory cytokines, growth factors, and metalloproteases, which constitute the senescence-associated secretory phenotype (SASP)^3, 8^.

PKCθ is a serine/threonine kinase that is well known for its role in the immune system as a key enzyme in T lymphocyte activation^9, 10^. Consequently, it is implicated in the T-cell−mediated diseases^9^. However, increasing evidence indicates that PKCθ is also involved in the pathology of cancer^11^. Indeed, PKCθ is highly expressed in estrogen receptor-negative breast tumors including TNBC cells and also in gastrointestinal stromal tumor (GIST), and it plays an active role in the development and progression of these types of cancer^12–20^. We and others have previously found that the PKCθ inhibition reduces the proliferation ability of breast cancer cells and GIST cells^14, 21, 22^. However, the function of PKCθ in cell proliferation and its related mechanism remain poorly understood.

Here we demonstrate that the PKCθ inhibition leads to a growth arrest combined with features of senescence in TNBC cells. Mechanistically, PKCθ silencing triggers senescence in a p53-independent manner through the downregulation of GADD45a with the consequent accumulation of p27.

## Methods

### Cell culture

Triple negative breast cancer cell lines: MDA-MB-436, MDA-MB-231 and HCC1937 were purchased from ATCC. These three breast cancer cell lines were cultured in medium according to ATCC’s recommendation. All cell lines were authenticated by Eurofins Genomics GmbH and were routinely tested for mycoplasma.

### siRNA transfection

Cells are transfected with siRNA duplexes (3,6 nM) by reverse transfection using interferin (Polyplus) according to manufacturer’s instructions. All siRNA duplexes were synthesized by Eurofins Genomics GmbH. The sequences of the siRNAs are:

siCtrl: 5’- AGGTAGTGTAATCGCCTTG-3’

siPKCθ-1 is a pool of 2 siRNAs: 5’-GACCATCTCTGCAGATTAA-3’; 5’- AGTAGAGCTCCACGGTGGT-3’

siPKCθ-2: 5’-GCACTTGGAAAGGTTTCAA-3’

siGadd45ɑ is a pool of 3 siRNAs: 5’-GGAGGAAGTGCTCAGCAAA-3’; 5’- AGTCGCTACATGGATCAAT-3’; 5’-GTGCTCAGCAAAGCCCTGA-3’

sip27 is a pool of 2 siRNAs: 5’-GCAACCGACGATTCTTCTA-3’; 5’-GCGTTGGATGTA GCATTAT-3’

MDA-MB-436 and MDA-MB-231 are transfected twice with siPKCθ-1 (72 hours between each transfection), and HCC1937 are only transfected once, in order to achieve the knockdown efficiency.

### Clonogenic assay

Cells are transfected with siRNA as described in transfection section. MDA-MB-436 and MDA-MB-231: 1,5.10^5^ cells are seeded in 6-well plate directly after the second transfection. HCC1937: 1,5.10^5^ cells are seeded in 6-well plate directly after the first transfection. Ten days later the cells are fixed with paraformaldehyde 4% during 10 minutes at room temperature then stained with crystal violet for 30 minutes protected from light. Plates dry overnight at room temperature. Images are taken by Chemidoc Imaging System and quantification is realized with ImageJ software: binary filter is applied then total number of black pixels is determined (value =255) and set as reference for control.

### EdU assay

Cells are transfected with siRNAs as described in transfection section: 5.10^3^ transfected cells are plated in 96 well-plates coated with 1µg/µL of fibronectin for 30 minutes. EdU (5 µM) is added to the medium for 1-2 hours (cell cycle analysis) or 24-48 hours (cell cycle arrest) prior fixation. Cells are fixed with paraformaldehyde 4% during 15 minutes, permeabilized with 1% Triton/PBS for 10 minutes and saturation was done with 1% BSA/PBS for 30 minutes. Cells are stained with Click-iT EdU Cell proliferation kit for Imaging (Alexa Fluor 647 dye) according to manufacturer’s instructions. 36 fields photos from each well (>500 cells) are taken by a confocal microscope Operetta –High Content Imaging System PERKIN ELMER (x20). Data analysis is done with Colombus software: the sum of DAPI and EdU intensity are measured individually for each cell. For cell cycle analysis (Figure 1D), cells are plotted on a graph by DAPI Sum (x-axis) and EdU sum intensity (y-axis) (Supplementary Figure 1B). Cell cycle phase is manually determined by the ratio of DAPI to EdU: G1 (Low DAPI/Low EdU), S (High DAPI/High EdU), G2/M(High DAPI/Low EdU). For cell cycle arrest, EdU positive cells are defined by EdU sum intensity and then quantified as percentages.

**Figure 1.**
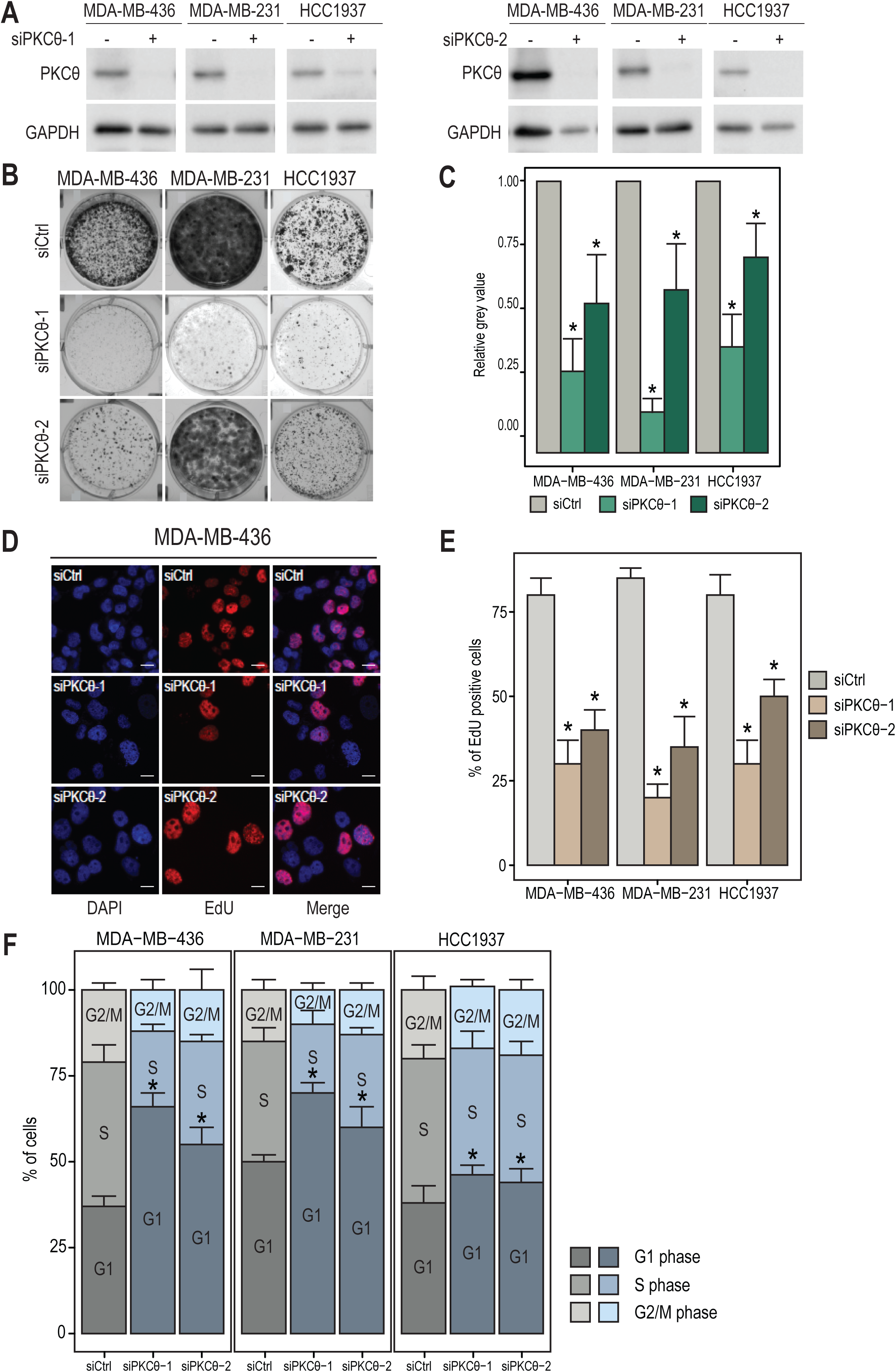
PKCθ loss leads to proliferation arrest in G1 phase. (A) Whole Cell Extracts (WCEs) from the indicated TNBC cells, transfected with two different siRNA targeting *PRKCQ* gene (siPKCθ-1 and siPKCθ-2), were analyzed by immunoblotting using antibodies against PKCθ and GAPDH. (B) The indicated TNBC cells, transfected with two different siRNA targeting *PRKCQ* gene, were subjected to a clonogenic assay for 14 days. Images from one representative assay (n= 3 independent experiments). (C) Quantification of images from clonogenic assays using ImageJ. Mean value of grey value +/- standard deviation from 3 independent experiments (control = 1). (D) MDA-MB-436 cells, transfected with two different siRNA targeting *PRKCQ* gene, were subjected to 24h EdU incorporation and DAPI staining. Images of DAPI and EdU staining from one representative assay (n= 3 independent experiments). scale bar = 15 µm. (E) The indicated TNBC cells, transfected with two different siRNA targeting *PRKCQ* gene, were subjected to 24h or 48h EdU incorporation and then the EdU positive cells were quantified (Mean percentage value of stained cells +/- standard deviation from 3 independent experiments). (F) The indicated TNBC cells, transfected with two different siRNA targeting *PRKCQ* gene, were subjected to 1h or 2h EdU incorporation and DAPI staining. The percentage of cells in each cell cycle phase was then quantified using Columbus software (n= 3 independent experiments). C, E, F: Significant differences are indicated with asterisks (*: p value < 0.01).

### Betagalactosidase assay

Cells are transfected as described in transfection section then seeded in 12-wells plates. 72h hours later, cells are fixed and marked following manufacturer’s instructions BioVision “Senescence detection Kit”. Images are acquired by Chemidoc Imaging System. Quantifications is manually done from pictures. An independent experience is done in triplicate with at least 100 cells counted in each triplicate. Results are the representation of mean +/- standard deviation from three independent experiences.

### Irradiation

Cells are plated in 96 wells and the next day cells are irradiated (5 Gy or 15 Gy) or mock-exposed (0 Gy) with a Cs^137^ source (Biobeam 8000). After 5 days of incubation, cells are fixed and stained with DAPI.

### Western Blot

Cells are scrapped at 4°C in lysis buffer (10mM EDTA, 5mM MgCl_2,_ 1% Triton X-100, 40 mM Tris pH8, protease/phosphatase inhibitors). 20µg of cell extracts are used for immunoblotting. Antibodies are incubated for the night at 4°C. Images are taken by Chemidoc Imaging System. Quantification is realized with Image Lab Software 5.2.1 (Biorad). The PKCθ, p21 and p27 antibodies were ordered from Cell Signalling Technologies. The p16, GADD45a and GAPDH antibodies were from Abcam, Millipore, and GeneTex, respectively.

### Transcript level analysis

Total RNA was extracted using TRIzol Reagent (Invitrogen). RT-qPCR analysis was performed using Takyon™ No ROX SYBR 2X MasterMix blue dTTP (Eurogentec). The primers used for transcriptional analysis are listed in Supplementary Table.

### PCR microarrays

RT^2^ Profiler PCR Arrays Human Cellular Senescence (PAHS-050ZD-2) from Qiagen were used to identify the gene expression profiles of MDA-MB-436 cells expressing siCtrl or siPKCθ-1. The arrays were performed twice in triplicate for each condition according to the manufacturer’s instructions.

### Statistics

All data are presented as mean +/- standard deviation. Statistical analysis to compare results among groups was carried out by the Student’s t test (GraphPad Prism software or excel software). A value of p < 0.05 was considered to be statistically significant.

## Results

### PKCθ loss leads to a proliferation arrest in G1 phase

To investigate the role of PKCθ in cell proliferation, we inhibited the PKCθ expression with two different siRNAs in several TNBC cells that express high level of PKCθ (Fig. 1A). The clonogenic cell survival assays, performed in MDA-MB-436, MDA-MB-231 and HCC1937 cells, showed that the PKCθ silencing suppressed the TNBC cell growth (Fig. 1B and C). We confirmed this observation with an EdU assay (Fig. 1D), thereby demonstrating that the PKCθ silencing reduced the proliferation rate of TNBC cells (Fig. 1E). Then, we asked in which cell cycle phase the TNBC cells were blocked. By combining DAPI and EdU fluorescent intensity (Suppl Fig. 1), we found an increased number of cells in G1 phase upon the PKCθ knockdown (Fig. 1F). Altogether, these data shows that the PKCθ loss in TNBC cells leads to a proliferation arrest in G1.

### PKCθ loss induces a cellular senescence-like phenotype

The proliferation arrest in G1 phase is often linked with the cell senescence^3, 8^. Thus, we verified whether the observed proliferation arrest was due to cell senescence. Here, we examined different senescence-like features^3, 8^. The PKCθ silencing in the TNBC cells led to an increase in the number of cells staining positive for SA-β-Gal activity (Fig. 2A-B). The proliferation arrest induced by the PKCθ loss was also accompanied by a senescence-associated secretory phenotype. Indeed, the PKCθ silencing caused an increase in mRNA levels of some chemokines (such as IL-1β, IL-8, IL-1α and CXCL2, but not IL-6) in the indicated breast cancer cell lines (Fig. 2C). However, we did not detect any SAHF formation upon the PKCθ loss, and even after irradiation which normally induces SAHF^23^, suggesting that our studied cancer cells were not able to form SAHF (Fig. 2D and Suppl Fig. 2). The senescence response is executed mainly through the activation of growth suppressors, including p53 and the cyclin-dependent kinase (CDK) inhibitors p21, p16 and p27. All studied TNBC cell lines harbor a loss of function mutation for p53^24^. Although MDA-MB-231 cells harbor a loss of p16 due to a mutation^25^ and a lack of p21 expression, we detected a significant increase of p21 expression and p16 expression at mRNA levels in MDA-MB-436 and HCC1937 cells upon PKCθ silencing while only p21 expression was enhanced at protein level (Fig. 3A-B). Importantly, PKCθ loss led to a prominent increase in p27 expression at both protein and mRNA levels in all three TNBC cell lines (Fig. 3 A-B). Thus, our results demonstrate that PKCθ silencing drives the TNBC cells into a senescence-like phenotype.

**Figure 2.**
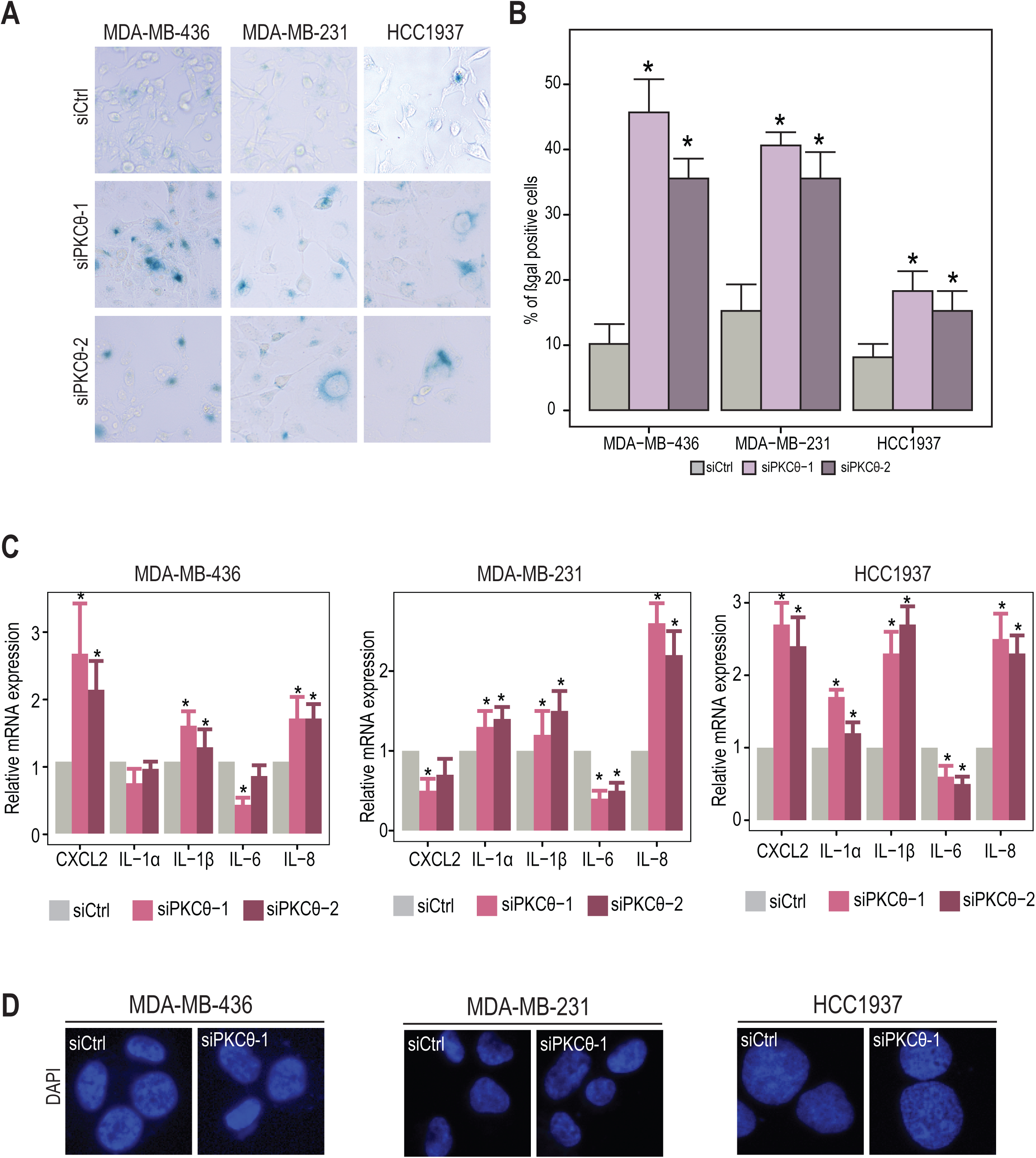
PKCθ loss induces a cellular senescence-like phenotype. (A) The indicated TNBC cells, transfected with two different siRNA targeting *PRKCQ* gene, were subjected to a β-galactosidase assay. Images from one representative assay (n= 3 independent experiments). (B) The indicated TNBC cells, transfected with two different siRNA targeting *PRKCQ* gene, were subjected to a β-galactosidase assay and then the β-galactosidase positive cells were quantified (n= 3 independent experiments). (C) The indicated TNBC cells were stained with DAPI to show the absence of SAHF formation. (D) Total RNA, extracted from the indicated TNBC cells transfected with two different siRNA targeting *PRKCQ* gene, was analyzed for the mRNA expression of the indicated genes (n= 3 independent experiments). B, C: Significant differences are indicated with asterisks (*: p value < 0.01).

**Figure 3.**
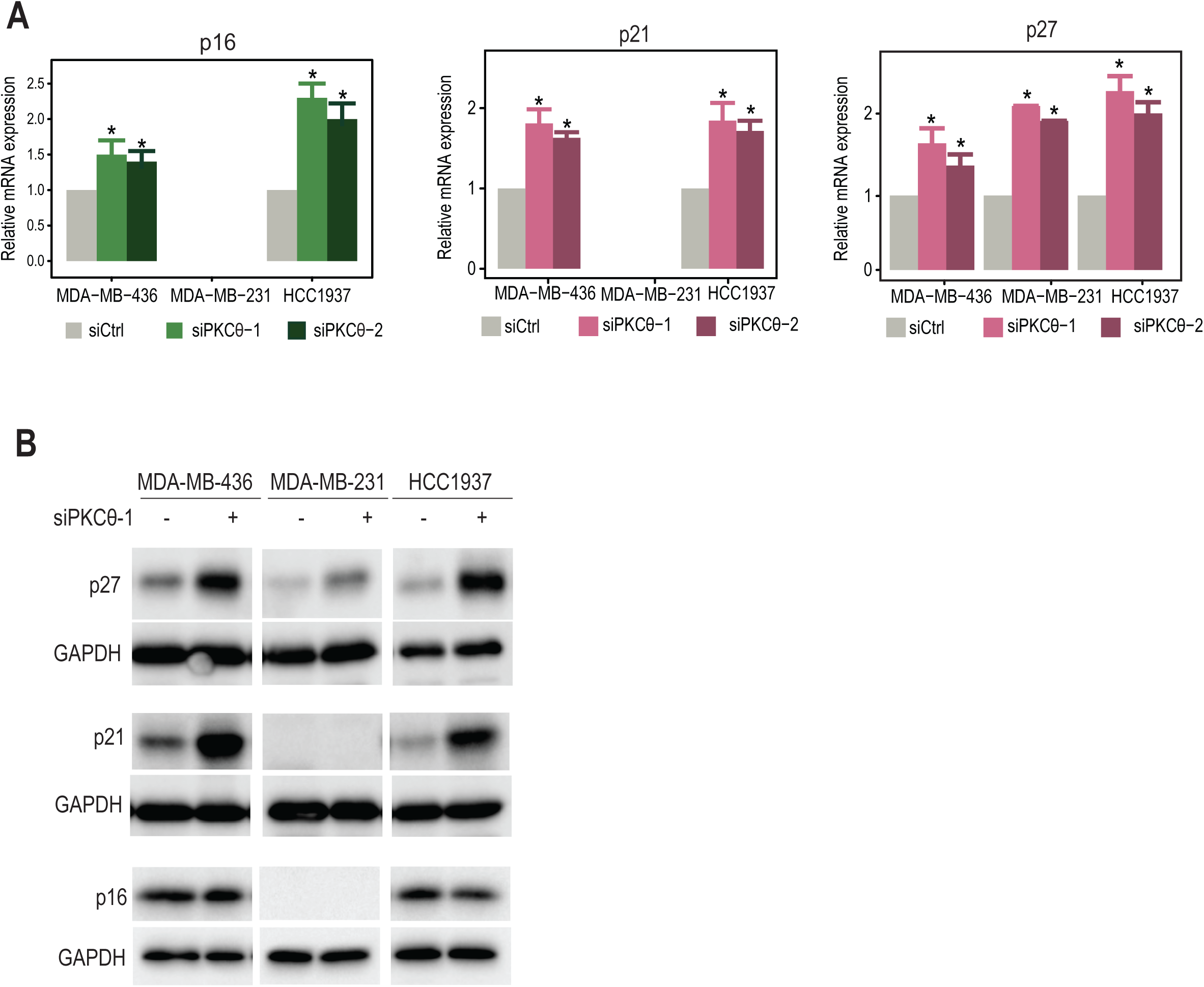
PKCθ loss strongly induces p27 expression in TNBC cells. (A) Total RNA, extracted from the indicated TNBC cells transfected with two different siRNA targeting *PRKCQ* gene, was analyzed for the mRNA expression of the indicated genes (n= 3 independent experiments). Significant differences are indicated with asterisks (*: p value < 0.01). (B) Whole-cell extracts (WCEs) from the indicated TNBC cells expressing siCtrl (-) or siPKCθ (+) were analyzed by immunoblotting using antibodies against PKCθ, p27, p21, p16 and GAPDH (n= 3 independent experiments).

### p27 inhibition rescues the phenotype induced by PKCθ depletion

Our data suggests that p27 is the common driver of PKCθ loss-induced senescence in the TNBC cell lines. To test whether p27 accumulation actively participates in the senescence induction, we silenced p27 in MDA-MB-436 and MDA-MB-231 cells concomitantly with PKCθ silencing (Fig. 4A). Preventing the accumulation of p27 significantly, while not completely, hampered the proliferation arrest induced by PKCθ loss as measured by the clonogenic and EdU assays (Fig. 4B-D), and blocked the increase in the number of cells staining positive for SA-βgal activity (Fig. 4E). Therefore, our results implicate p27 as the main molecular target driving the PKCθ loss-induced senescence in TNBC cells.

**Figure 4.**
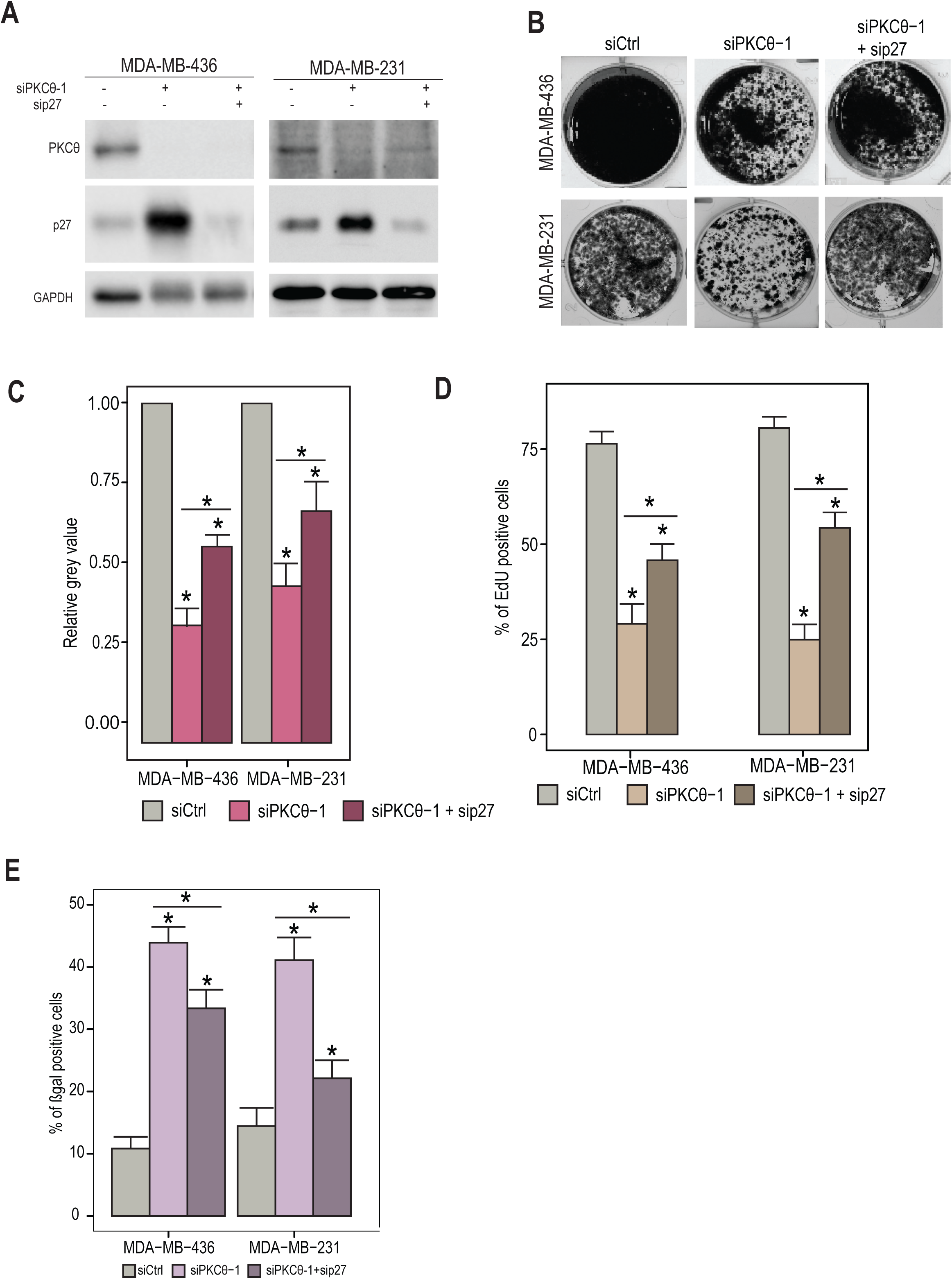
p27 silencing rescues the phenotype induced by PKCθ loss. (A) WCEs from the indicated TNBC cells expressing siCtrl (-) or siPKCθ (+) and/or sip27 (+) were analyzed by immunoblotting using antibodies against PKCθ, p27and GAPDH. (B) The indicated TNBC cells, transfected with siPKCθ-1 and/or sip27, were subjected to a clonogenic assay for 14 days. Images from one representative assay (n= 3 independent experiments). (C) Quantification of images from clonogenic assays using ImageJ. Mean value of grey value +/- standard deviation from 3 independent experiments (control = 1). (D) The indicated TNBC cells, transfected with siPKCθ-1 and/or sip27, were subjected to 24h EdU incorporation and then the EdU positive cells were quantified (Mean percentage value of stained cells +/- standard deviation from 3 independent experiments). (E) The indicated TNBC cells, transfected with siPKCθ-1 and/or sip27, were subjected to a β-galactosidase assay and then the β-galactosidase positive cells were quantified (n= 3 independent experiments). C, D, E: Significant differences are indicated with asterisks (*: p value < 0.01).

### PKCθ loss causes the dysregulation of various genes involved in cell proliferation and senescence

To better decipher the molecular pathway driving the PKCθ loss-induced senescence, we performed expression analysis of genes known to regulate cell proliferation and senescence in MDA-MB-436 cells (Table 1). Overall, the PKCθ loss was accompanied by the up-regulation and down-regulation of various genes implicated in cell proliferation and senescence, further supporting that the activation of a senescence-like phenotype is associated with striking epigenetic changes. As expected, we found a significant induction in CDKN2A (p16), CDKN1B (p27), CDKN1A (p21) and GLB1 (SA-βgal) expression following the PKCθ silencing. Consistent with cell proliferation arrest and senescence, PKCθ loss induced a decrease in MYC and PCNA expression while an increase in ATM and CREG1 expression, which were confirmed in MDA-MB-436, MDA-MB-231 and HCC1937 cells (Suppl Fig 3). Surprisingly, this analysis highlighted a strong down-regulation of GADD45a (Growth Arrest and DNA Damage Inducible Alpha) mRNA level. GADD45a expression, known to be rapidly and strongly induced in response to cellular stress, usually results in cell cycle arrest and DNA repair^26, 27^. However, we observed the strong GADD45a reduction during TNBC cell senescence. To further validate this unexpected result, we checked the expression of GADD45a following the PKCθ inhibition in all TNBC cell lines. Consistent with the gene expression analysis, PKCθ loss resulted in the strong down-regulation of GADD45a at both mRNA (Fig. 5A) and protein (Fig. 5B) levels. Thus, the gene expression analysis reveals that PKCθ loss dysregulates the expression of a significant number of genes involved in cell proliferation and senescence, especially a surprising decrease in GADD45a level.

**Figure 5.**
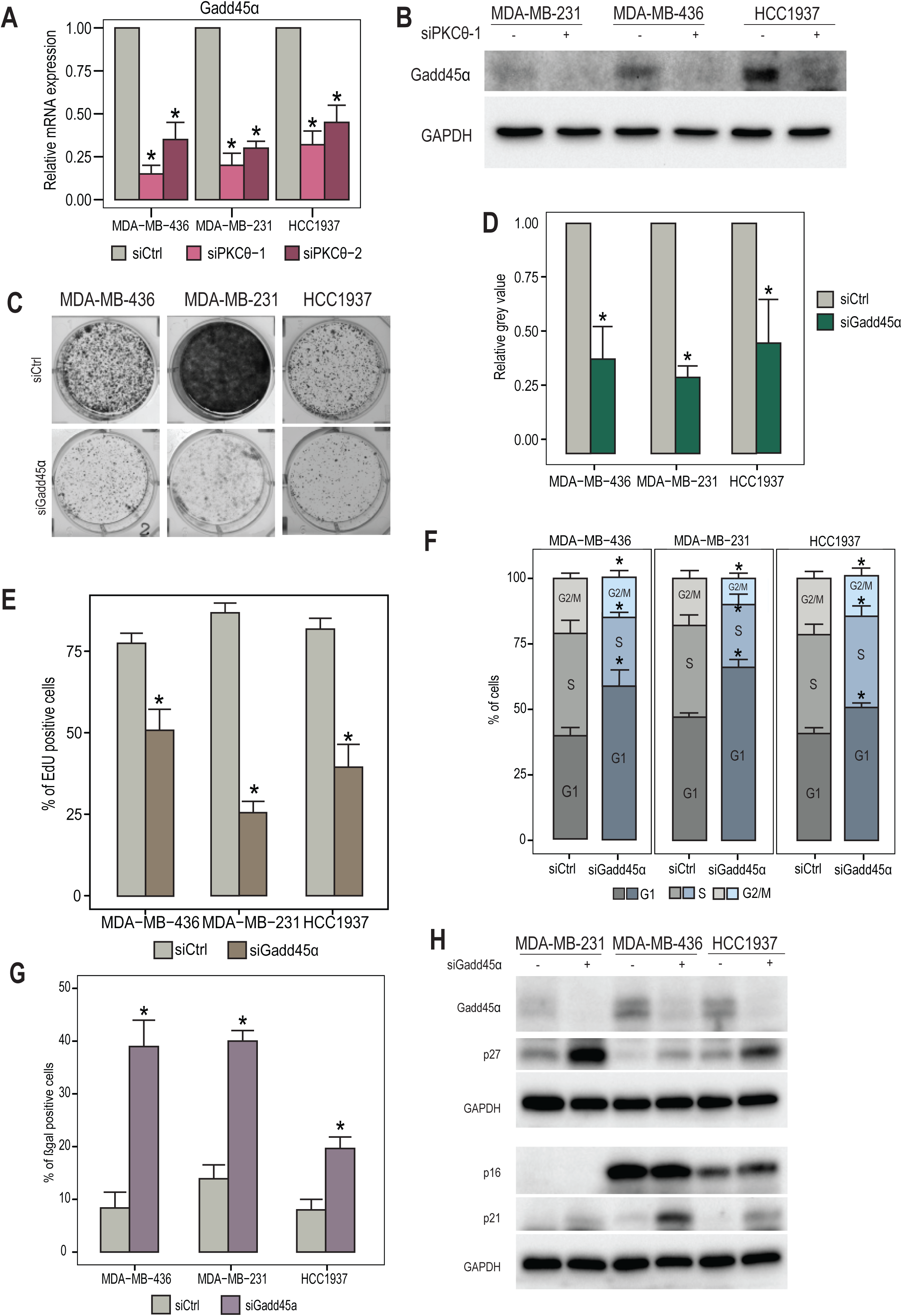
GADD45a silencing leads to a senescence-like phenotype. (A) Total RNA, extracted from the indicated TNBC cells transfected with two different siRNA targeting *PRKCQ* gene, was analyzed for the mRNA expression of the *GADD45a* gene (n= 3 independent experiments). (B) WCEs from the indicated TNBC cells expressing siCtrl (-) or siPKCθ (+) were analyzed by immunoblotting using antibodies against GADD45a and GAPDH (n= 3 independent experiments). (C) The indicated TNBC cells, transfected with a pool of three different siRNAs targeting *GADD45a* gene (siGADD45a), were subjected to a clonogenic assay for 14 days. Images from one representative assay (n= 3 independent experiments). (D) Quantification of images from clonogenic assays using ImageJ. Mean value of grey value +/- standard deviation from 3 independent experiments (control = 1). (E) The indicated TNBC cells, transfected with siGADD45a, were subjected to 24h EdU incorporation and then the EdU positive cells were quantified (Mean percentage value of stained cells +/- standard deviation from 3 independent experiments). (F) The indicated TNBC cells, transfected with siGADD45a, were subjected to 1h or 2h EdU incorporation and DAPI staining. The percentage of cells in each cell cycle phase was then quantified using Columbus software (n= 3 independent experiments). (G) The indicated TNBC cells, transfected with siGADD45a, were subjected to a β-galactosidase assay and then the β-galactosidase positive cells were quantified (n= 3 independent experiments). (H) WCEs from the indicated TNBC cells expressing siCtrl (-) or siGADD45a (+) were analyzed by immunoblotting using antibodies against GADD45a, p16, p21, p27 and GAPDH (n= 3 independent experiments). D, E, F, G: Significant differences are indicated with asterisks (*: p value < 0.01).

**Table 1.** Expression analysis of genes involved in cell proliferation and senescence. MDA-MB-436 cells expressing siCtrl or siPKCθ were analyzed using the human cellular senescence PCR array profiler. mRNA levels of the indicated genes were quantified by RT-QPCR following PKCθ silencing. The table shows the results of 2 independent experiments.

| Genes | Exp#1 | Exp#2 | Genes | Exp#1 | Exp#2 | Genes | Exp#1 | Exp#2 |
| --- | --- | --- | --- | --- | --- | --- | --- | --- |
| CREG1 | 2.65 | 2.09 | SOD2 | 1.12 | 1.05 | NBN | 2.26 | 1.68 |
| CDK6 | 2.17 | 1.9 | COL1A1 | 1.18 | 1.32 | CDKN2D | 2.34 | 1.64 |
| TGFB1 | 2.08 | 1.87 | SERPINB2 | 1.17 | 1.44 | RBL1 | 2.38 | 1.33 |
| GLB1 | 2.07 | 1.85 | E2F3 | 1.25 | 1.01 | CCNB1 | 2.42 | 1.28 |
| CDKN2A | 1.97 | 1.5 | SPARC | 1.25 | 1.42 | CDK2 | 2.46 | 1.67 |
| BMI1 | 1.71 | 1.75 | RB1 | 1.34 | 1.23 | CCND1 | 2.54 | 1.11 |
| CDKN1B | 1.55 | 1.86 | AKT1 | 1.42 | 1.12 | ING1 | 2.68 | 1.68 |
| MAP2K3 | 1.45 | 1.15 | CDKN2C | 1.44 | 1.07 | SOD1 | 2.7 | 2.4 |
| ALDH1A3 | 1.4 | 1.7 | TGFB1I1 | 1.45 | 1.05 | CHEK1 | 2.99 | 2.25 |
| ID1 | 1.39 | 2.62 | PLAU | 1.5 | 1.27 | TP53 | 3.16 | 2.09 |
| CD44 | 1.33 | 1.54 | MORC3 | 1.52 | 1.67 | PTEN | 3.27 | 3 |
| CDKN1A | 1.31 | 1.02 | CDK4 | 1.6 | 1.27 | FN1 | 4.03 | 2.67 |
| IGFBP3 | 1.31 | 1.13 | EGR1 | 1.62 | 2.94 | CCNA2 | 4.57 | 1.95 |
| ATM | 1.27 | 1.31 | HRAS | 1.73 | 1.81 | PCNA | 4.66 | 2.52 |
| SERPINE1 | 1.27 | 1.86 | TP53BP1 | 1.73 | 1.52 | ETS1 | 4.93 | 3.18 |
| CDKN2B | 1.24 | 1.16 | IGF1R | 1.81 | 1.35 | MYC | 4.95 | 3.98 |
| MDM2 | 1.23 | 1 | IRF3 | 1.87 | 1.67 | E2F1 | 5.26 | 2.25 |
| PIK3CA | 1.2 | 1.14 | IRF5 | 1.9 | 1.77 | TBX3 | 6.35 | 2.62 |
| CALR | 1.19 | 1.27 | RBL2 | 1.88 | 1.56 | IRF7 | 6.98 | 2.34 |
| CCNE1 | 1.14 | 1.38 | SIRT1 | 1.88 | 1.72 | GADD45A | 15.38 | 4.54 |
| CITED2 | 1.13 | 1.73 | CDC25C | 1.94 | 1.07 | THBS1 | 13.63 | 14.4 |
| VIM | 1.12 | 1.06 | IGFBP7 | 1.91 | 1.84 | MAP2K6 | 235.43 | 31.85 |
| GSK3B | 1.01 | 1 | ETS2 | 2 | 1.94 | COL3A1 |  | 19.04 |
| ABL1 | 1.01 | 1.07 | TERF2 | 2.02 | 1.64 | IFNG |  |  |
| TWIST1 | 1.02 | 1.05 | MAP2K1 | 2.04 | 1.7 | IGF1 |  |  |
| PRKCD | 1.08 | 1.3 | MAPK14 | 2.1 | 1.69 | IGFBP5 |  |  |
| CHEK2 | 1.13 | 1.1 | TBX2 | 2.09 | 3.81 | NOX4 |  |  |
| NFKB1 | 1.13 | 1.33 | CDKN1C | 2.28 | 2.63 | TERT |  |  |

### GADD45a reduction drives PKCθ loss-induced senescence

Having established GADD45a as a downstream target of PKCθ, we next explored the implication of this target in the activation of senescence induced by PKCθ loss. To do this, we inhibited GADD45a with a pool of three different siRNAs in MDA-MB-436, MDA-MB-231 and HCC1937 cells. Similar to the phenotype of PKCθ silencing, GADD45a inhibition reduced the proliferation rate of TNBC cells, and these cells were blocked in G1 phase (Fig. 5C-F and Suppl Fig. 4A-B). Furthermore, GADD45a inhibition also led to an increase in the number of cells staining positive for SA-βgal activity (Fig. 5G and Suppl Fig. 4C) and to a prominent induction of p27 and p21 protein expression while p16 protein increase was undetectable (Fig. 5H). Thus, our data demonstrate that GADD45a silencing drives the TNBC cells into a senescence-like phenotype. Importantly, the re-expression of GADD45a hampered the proliferation arrest induced by PKCθ loss as measured by the clonogenic and EdU assays (Fig 6A-E). GADD45a re-expression also partially blocked the increase in the number of cells staining positive for SA-βgal activity (Fig. 6F). Altogether, our data demonstrates that GADD45a strong reduction acts as a downstream effector of PKCθ loss to induce a senescence-like phenotype in TNBC cells.

**Figure 6.**
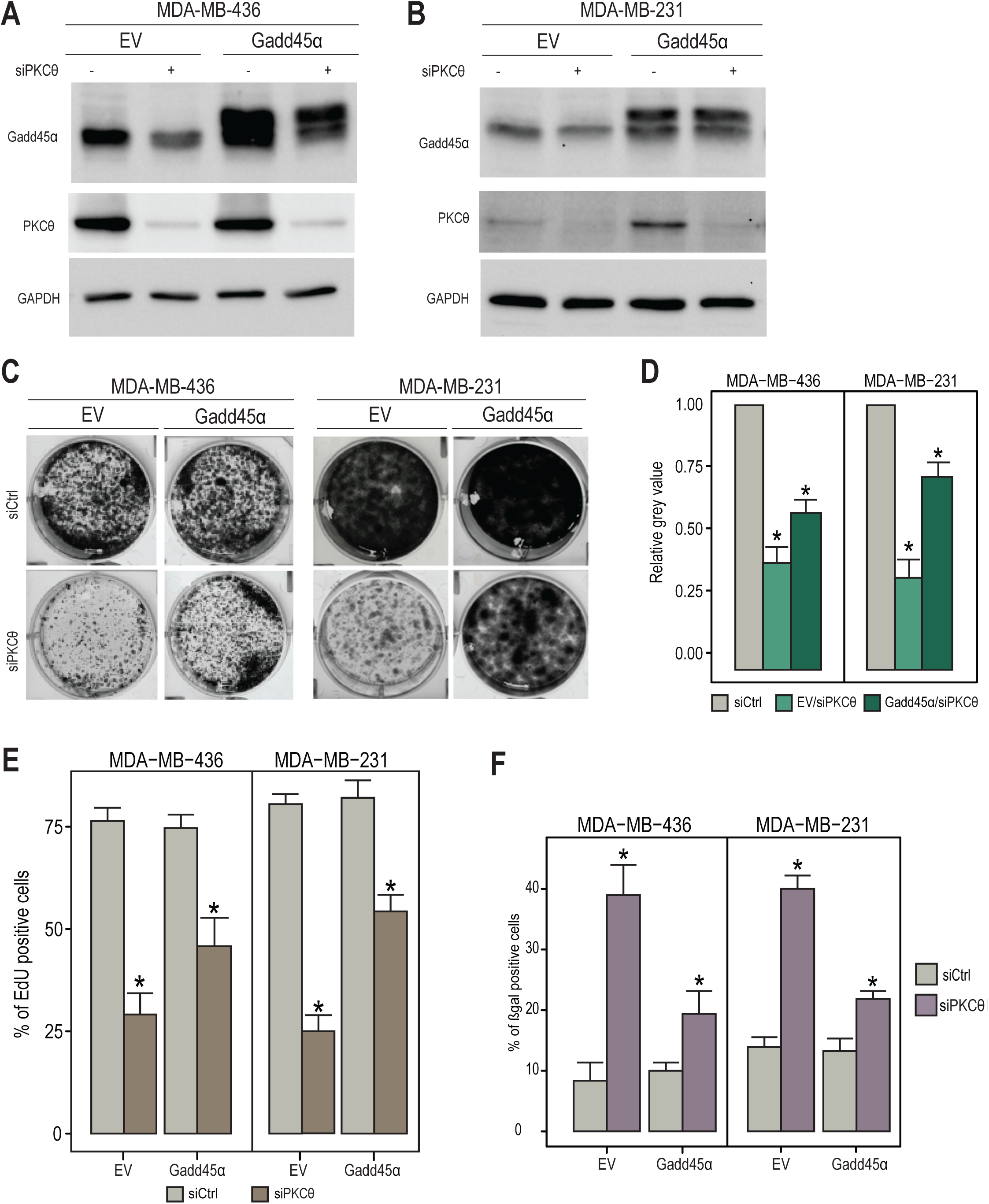
GADD45 down-regulation is responsible for the PKCθ loss-induced senescence. (A) The indicated TNBC cells stably expressing GADD45a were transfected with siCtrl (-) or siPKCθ (+). Two days later, WCEs were analyzed by immunoblotting using antibodies against GADD45a, PKCθ and GAPDH. (B) The indicated TNBC cells, stably expressing GADD45a and transfected with siCtrl or siPKCθ, were subjected to a clonogenic assay for 14 days. Images from one representative assay (n= 3 independent experiments). (C) Quantification of images from clonogenic assays using ImageJ. Mean value of grey value +/- standard deviation from 3 independent experiments (control = 1). (D) The indicated TNBC cells, stably expressing GADD45a and transfected with siCtrl or siPKCθ, were subjected to 24h EdU incorporation and then the EdU positive cells were quantified (Mean percentage value of stained cells +/- standard deviation from 3 independent experiments). (F) The indicated TNBC cells, stably expressing GADD45a and transfected with siCtrl or siPKCθ, were subjected to a β-galactosidase assay and then the β-galactosidase positive cells were quantified (n= 3 independent experiments). D, E, F: Significant differences are indicated with asterisks (*: p value < 0.01).

## Discussion

Here we found that PKCθ loss led to a growth arrest in TNBC cells. Consistently, we and others have previously reported that the inhibition of PKCθ by expression of a dominant negative form or by shRNA expression reduced the proliferation rate of breast cancer cells and GIST cells, and led to an increase of 27 expression ^14, 21, 22^. However, no senescence-like phenotype has been described in any of these cases. PKCθ regulates p27 expression through the Akt-FOXO3a pathway in MMTV-c-rel mouse mammary cells^14^, but this activation does not occur in TNBC cells (data not shown) suggesting that the response to PKCθ silencing might be different depending on the cellular context.

While the CDK inhibitors p16 and p21 are well known to trigger cell senescence, the accumulation of p27 has also been reported to participate in the activation of the senescence response^6, 7^. In our cellular context, p27 is the main driver of PKCθ loss-induced senescence. However, we also observed in some of the TNBC cells a slight induction of p16 proteins and an increase of p21 expression. Moreover, preventing the accumulation of p27 did not completely rescue cells from senescence, especially in MDA-MB-436 cells which express both p16 and p21. Altogether, these data suggest that for some TNBC cells, the senescence response might be triggered by the activation of several CDK inhibitors. Interestingly, the p21 induction is p53-independent in our cellular context. MYC has been reported to repress p21 expression and it could also be the case in our model since PKCθ silencing caused a decrease of MYC expression in TNBC cells^28–30^.

Our data showed that PKCθ loss-induced senescence is accompanied by a strong down-regulation of GADD45a and GADD45a silencing is also able to drive TNBC cells into a senescence-like phenotype. These results are somehow contradictory to the previously reported function of GADD45a since its up-regulation induced by various stresses is usually accompanied by a cell cycle arrest^31–33^. However, a few reports indicate that GADD45a is required for the survival of various types of cells^34–38^. In addition, GADD45a has been described as a tumor promoter or suppressor depending on the oncogenic stress in breast cancer^39, 40^. Indeed, while GADD45a promotes MYC-driven mammary tumorigenesis, it suppresses Ras-driven mammary tumorigenesis by regulating different signaling pathways^39–42^. Consistently, TNBC displays elevated levels and activity of diverse oncogenes including MYC, while Ras mutations that lead to constitutive activation of this cascade are not frequent in breast cancer^43^. Consistently, our data shows that PKCθ loss leads to a prominent downregulation of MYC expression in all TNBC cells. Interestingly, while GADD45a levels have been associated with estrogen receptor status, high GADD45a levels predicts a poor overall survival within the TNBC group^44, 45^. Therefore, GADD45a seems to behave as a tumor promoter in the context of TNBC cells.

GADD45a is an adaptor protein that binds to a variety of regulatory factors to modulate their function^27^. Depending on the stress stimuli, GADD45a interacts with PCNA, p21, histones, MEKK4, p38, MKK7 to promote epigenetic gene activation or kinase activation^26, 27^. GADD45a also acts as a heterochromatin relaxer by destabilizing histone–DNA interactions, thereby activating genes involved in somatic cell reprogramming^46^. In the context of TNBC cells, we observed a global increase in the trimethylation level of histone H3 Lysine-4 following the GADD45a silencing (data not shown, as well as PKCθ silencing). The Menin/MLL1 complex has been shown to regulate the expression of p27 and p16 genes by increasing the trimethylation of histone H3 Lysine-4 associated with their promoter^47, 48^. We could therefore envisage that GADD45 might modulate the methyltransferase activity of this complex to control the expression of the senescence drivers. Further studies would be needed to elucidate this mechanism.

The results obtained in this study further strengthen the novel oncogenic function of PKCθ in breast cancer^11, 20^. PKCθ is an attractive therapeutic target for TNBC, with expected minimal toxic side effects of specific PKCθ inhibitors since the PKCθ knock-out mice are generally healthy^49, 50^ and PKCθ expression is restricted to a few normal cell types^51^ while it is highly expressed in TNBC^13, 17, 20^. Targeting this kinase could be useful to reduce the aggressive behavior of TNBC cells by not only reducing its invasive ability but by also triggering a senescence response. Further studies are therefore required to quantify the level and activity of PKCθ in a large numbers of human breast cancer samples in order to validate this kinase as a therapeutic target for this aggressive subtype of breast cancers. In the case of GIST, PKCθ is used as a marker for the diagnosis of KIT-negative tumors^16^. Moreover, specific PKCθ inhibitors are currently being developed for therapeutic use and are undergoing preclinical testing in the context of T-cell-mediated diseases^52^. This clinical trial of PKCθ inhibitors could be also useful to treat aggressive breast cancer diseases.

Altogether, our data reveals the PKCθ-GADD45a axis as a novel regulatory mechanism preventing cancer cell senescence and further supports that targeting PKCθ can be beneficial to impair TNBC growth.

## Supporting information

Supplementary informations and Supplementary Figures

## Ethical Approval

Non applicable

## Availability of data and materials

All data generated and analyzed during this study are included in this published article. Materials generated in this study are available from the corresponding author upon reasonable request.

## ACKNOWLEDGEMENTS

We thank Malek Djabali and Maelle Carraz for helpful scientific discussion. We thank the LITC imaging facility. This work was supported by the INSERM Plan Cancer 2014-2019, La Ligue Contre le Cancer (Comité Haute-Garonne, 2019-2020), La Ligue National Contre le Cancer (PhD fellowship to A.N.) and Fondation ARC (PhD fellowship to A.N. and to Y.Z.), China Scholarship Council (PhD fellowship to Y.Z.)

## AUTHOR CONTRIBUTIONS

A.N. and K.B. designed the study. A.N., Z.Y., V.C. and K.B. conducted all experiments, analyzed data and interpreted results. A.N., Z.Y., X.W. and K.B. wrote the manuscript. K.B. supervised the overall project.

