## Supplementary informations and Supplementary Figures for "Loss of PKCθ-GADD45a axis drives triple-negative breast cancer cells into p53-independent senescence"

**Supplementary Figure legends**

**Supplementary Figure 1.** (A) The indicated TNBC cells, transfected with two different siRNA targeting *PRKCQ* gene (siPKCθ), were subjected to 24h or 48h EdU incorporation and DAPI staining. Images of DAPI and EdU staining from one representative assay (n= 3 independent experiments). scale bar = 15 µm. (B) The indicated TNBC cells, transfected with siPKCθ, were subjected to 1h or 2h EdU incorporation and DAPI staining. The data represents the intensity of Edu and DAPI staining (n= 3 independent experiments).

**Supplementary Figure 2.** MDA-MB-436 cells were irradiated and then stained with DAPI to show the absence of SAHF formation (n= 2 independent experiments).

**Supplementary Figure 3.** Total RNA, extracted from the indicated TNBC cells transfected with siPKCθ, was analyzed for the mRNA expression of the indicated genes (n= 3 independent experiments). Significant differences are indicated with asterisks (*: p value < 0.01).

**Supplementary Figure 4.** (A) The indicated TNBC cells, transfected with a pool of three different siRNA targeting *GADD45a* gene (siGADD45a), were subjected to 24h or 48h EdU incorporation and DAPI staining. Images of DAPI and EdU staining from one representative assay (n= 3 independent experiments). scale bar = 15 µm. (B) The indicated TNBC cells, transfected with siGADD45a, were subjected to 1h or 2h EdU incorporation and DAPI staining. The data represents the intensity of Edu and DAPI staining (n= 3 independent experiments). (C) The indicated TNBC cells, transfected with siGADD45a, were subjected to a β-galactosidase assay. Images from one representative assay (n= 3 independent experiments).

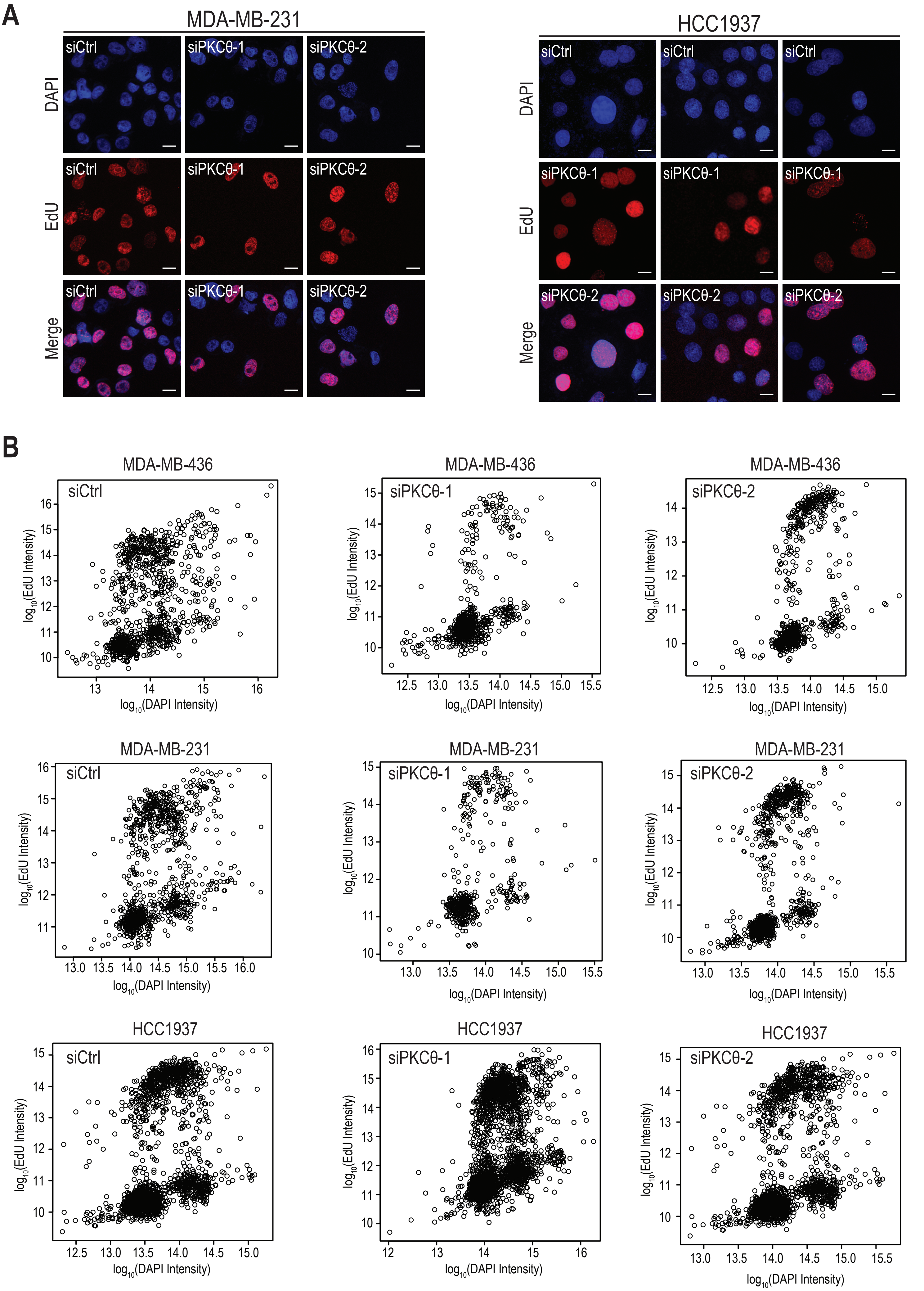


**Supplementary Figure 1**

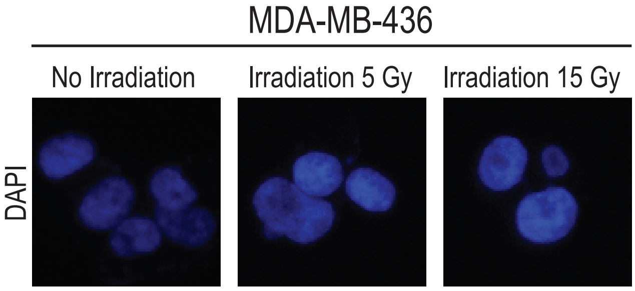


Supplementary Figure 2

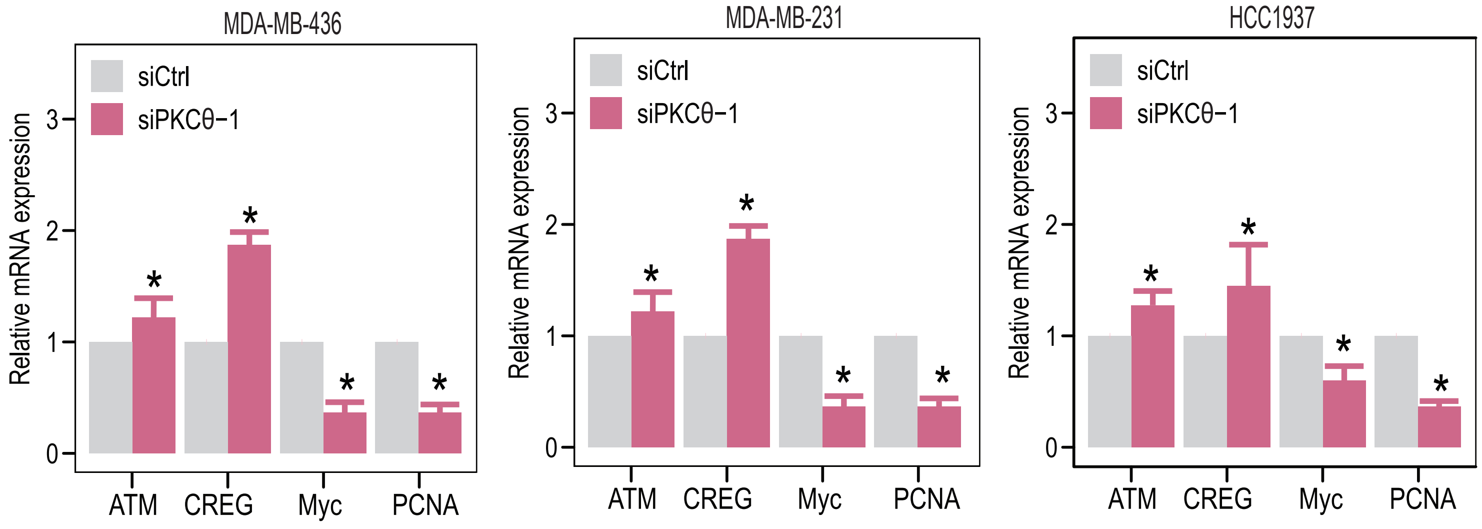


**Supplementary Figure 3**

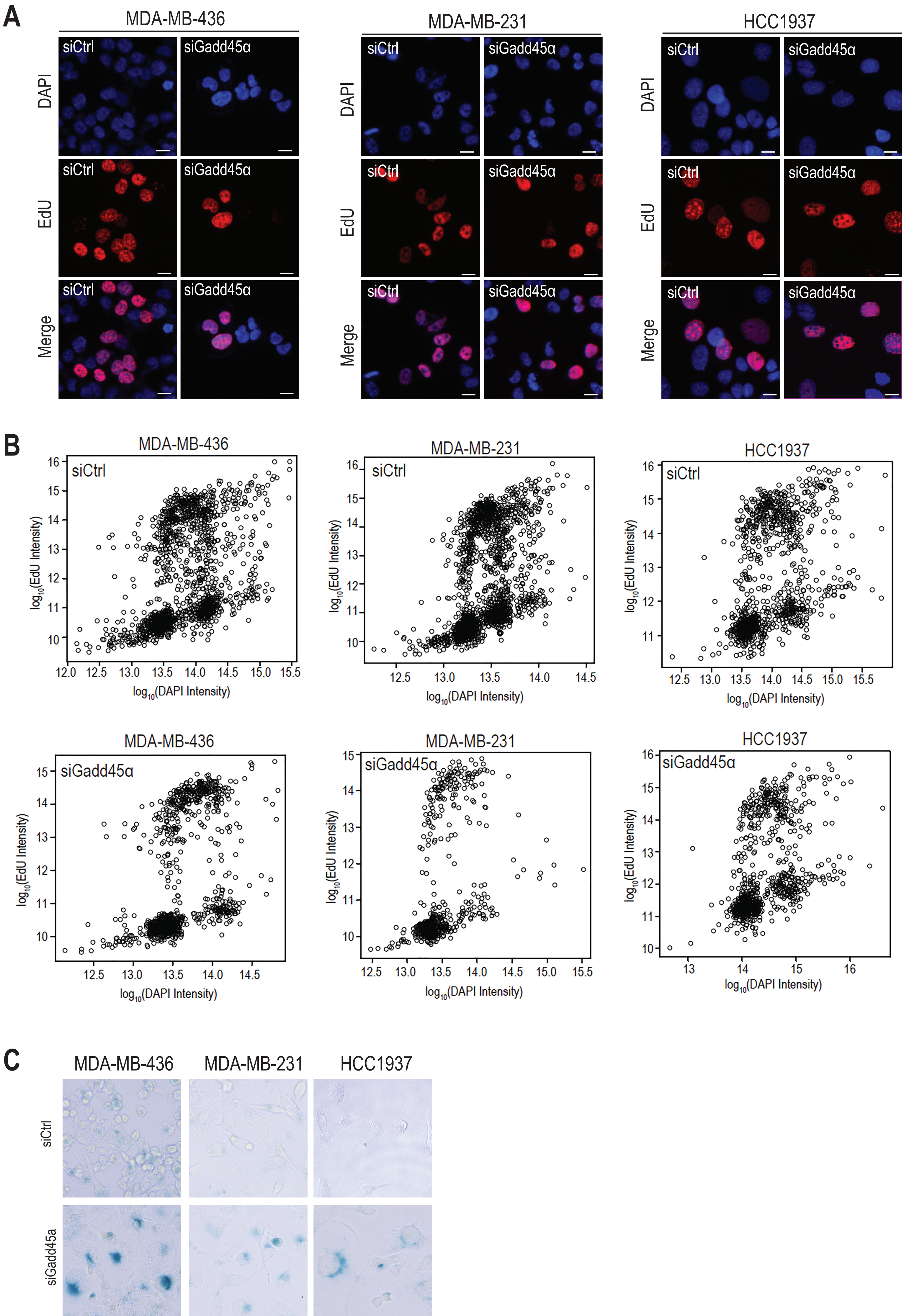


**Supplementary Figure 4**

Figure 1

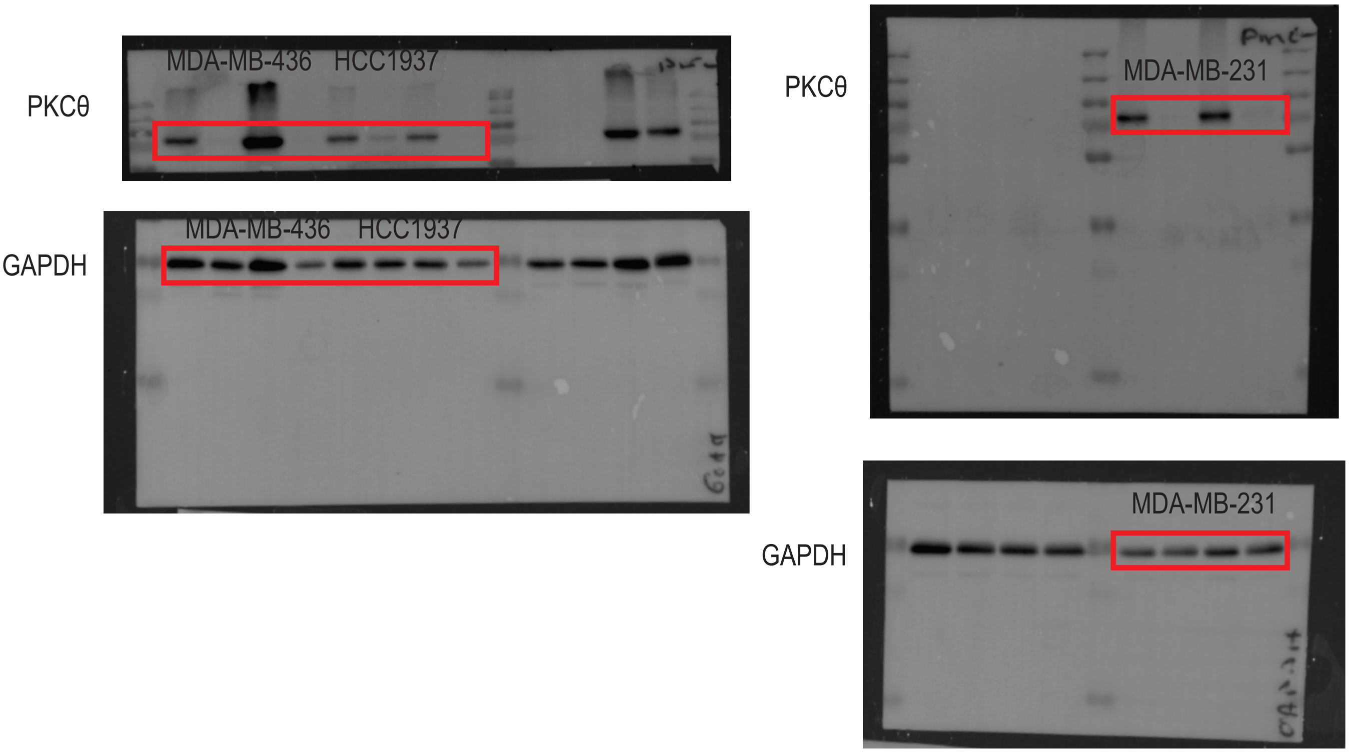


Figure 3

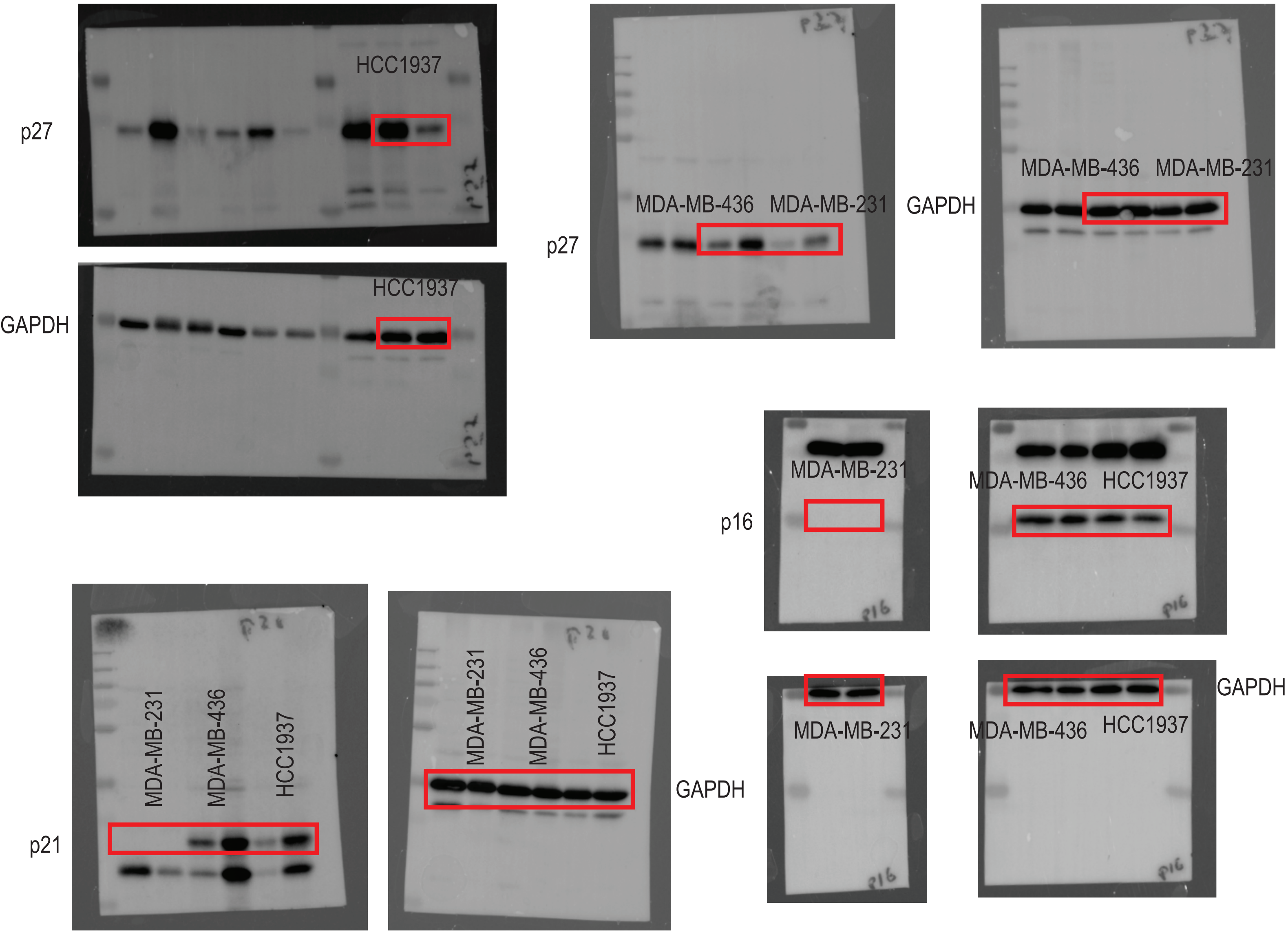


**Supplementary Figure 5**

Figure 4

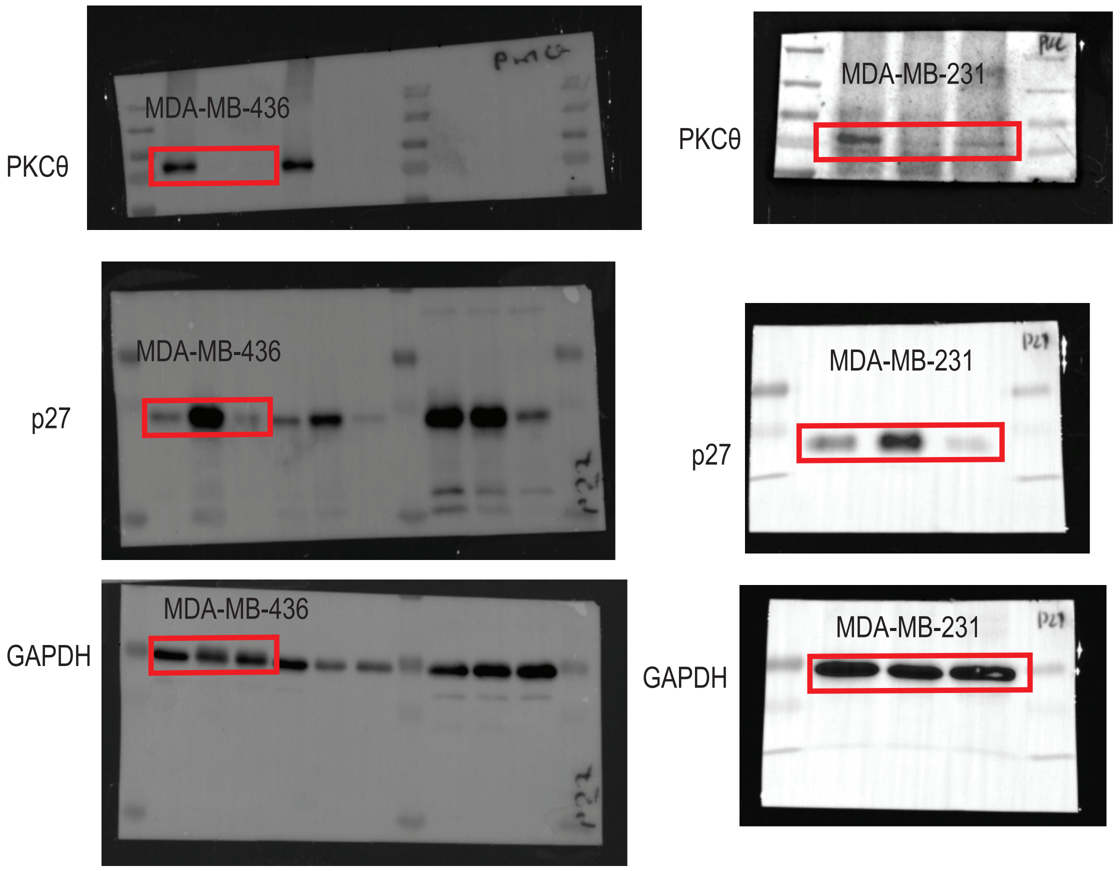


Figure 5B

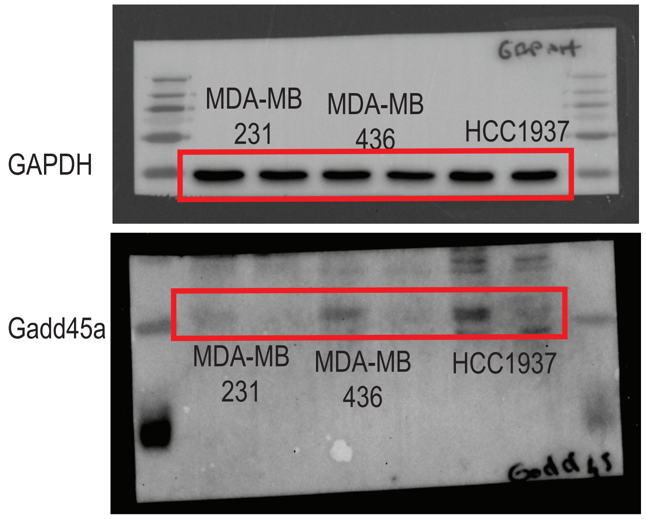


**Supplementary Figure 6**

Figure 5H

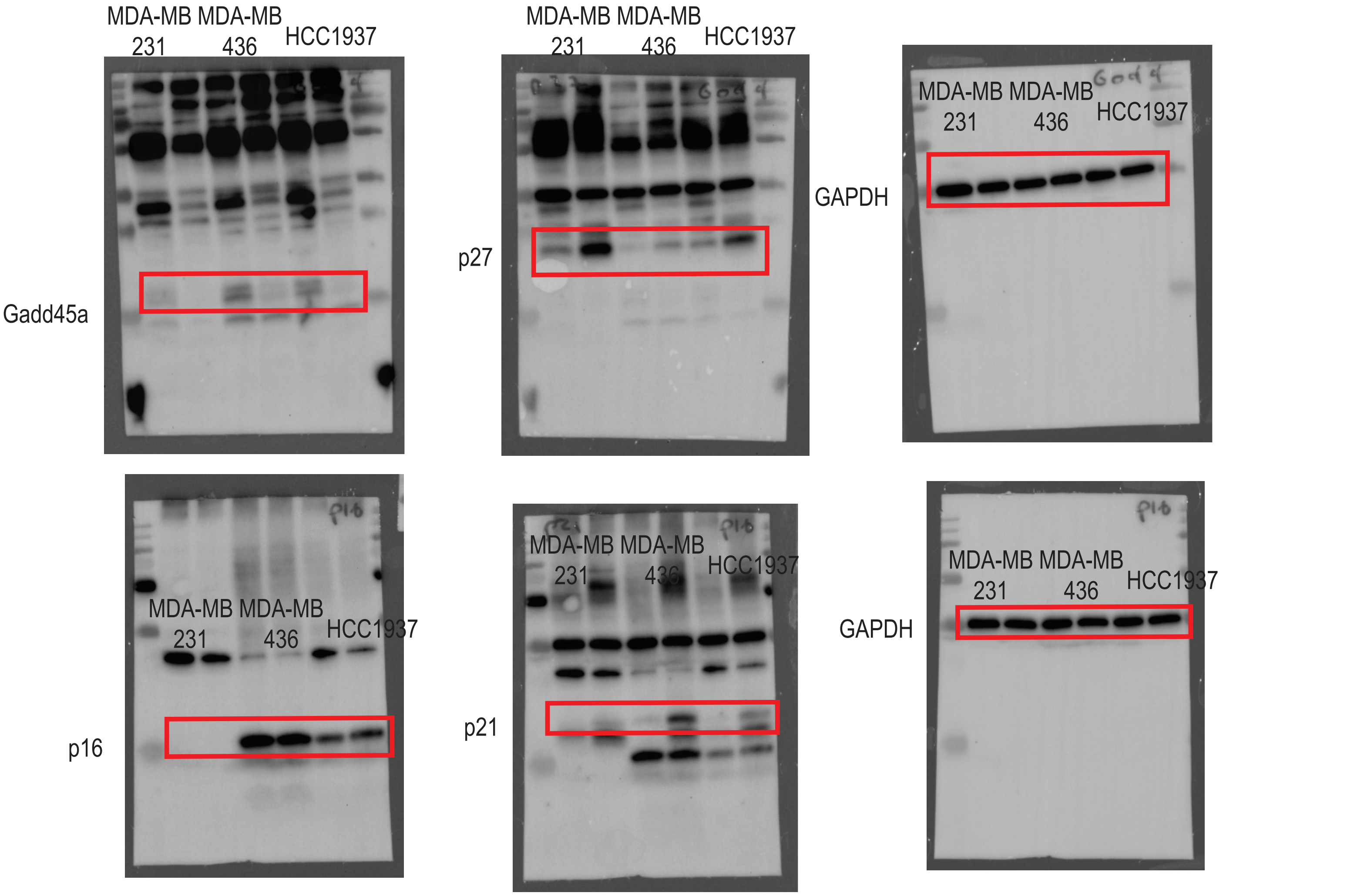


Figure 6

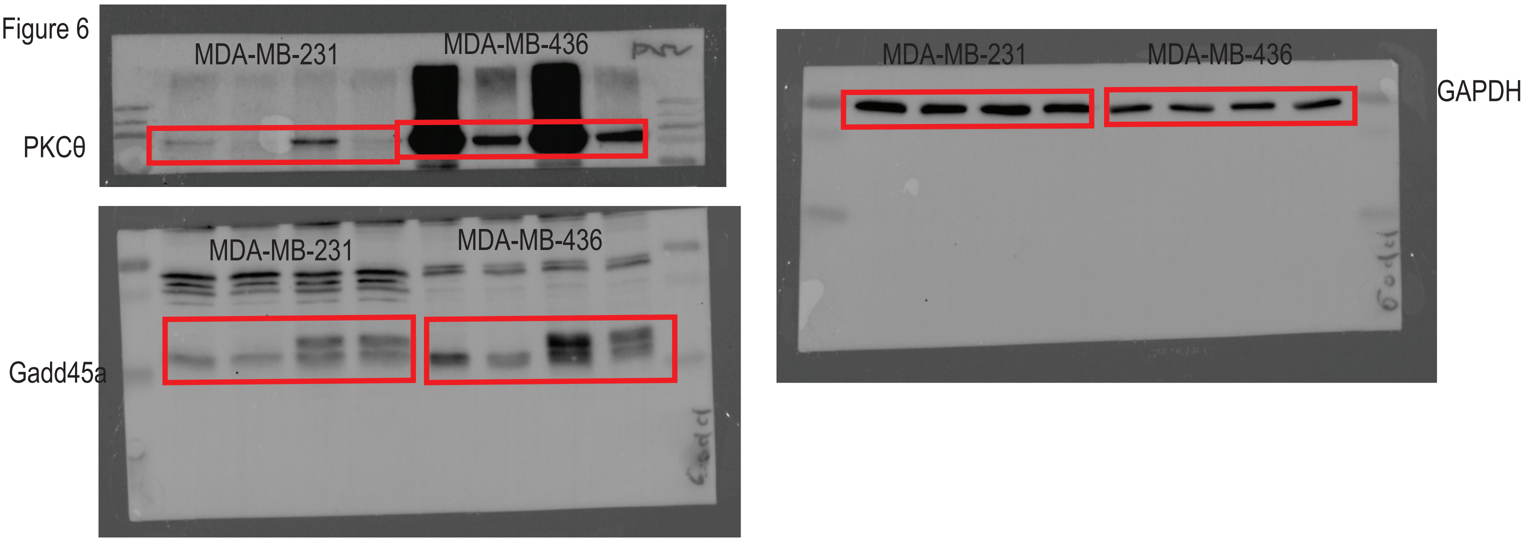


Supplementary Figure 7
